# A genomic approach to inferring kinship reveals limited intergenerational dispersal in the yellow fever mosquito

**DOI:** 10.1101/636456

**Authors:** Moshe Jasper, Thomas L. Schmidt, Nazni W. Ahmad, Steven P. Sinkins, Ary A Hoffmann

## Abstract

Understanding past dispersal and breeding events can provide insight into ecology and evolution, and can help inform strategies for conservation and the control of pest species. However, parent-offspring dispersal can be difficult to investigate in rare species and in small pest species such as mosquitoes. Here we develop a methodology for estimating parent-offspring dispersal from the spatial distribution of close kin, using pairwise kinship estimates derived from genome-wide single nucleotide polymorphisms (SNPs). SNPs were scored in 162 *Aedes aegypti* (yellow fever mosquito) collected from eight close-set, high-rise apartment buildings in an area of Malaysia with high dengue incidence. We used the SNPs to reconstruct kinship groups across three orders of kinship. We transformed the geographical distances between all kin pairs within each kinship category into axial standard deviations of these distances, then decomposed these into components representing past dispersal events. From these components, we isolated the axial standard deviation of parent-offspring dispersal, and estimated neighbourhood area (129 m), median parent-offspring dispersal distance (75 m), and oviposition dispersal radius within a gonotrophic cycle (36 m). We also analysed genetic structure using distance-based redundancy analysis and linear regression, finding isolation by distance both within and between buildings and estimating neighbourhood size at 268 individuals. These findings indicate the scale required to suppress local outbreaks of arboviral disease and to target releases of modified mosquitoes for mosquito and disease control. Our methodology is readily implementable for studies of other species, including pests and species of conservation significance.

## Introduction

Dispersal is a key trait in ecology and evolution, allowing species to evade stressful areas and locate favourable new areas, and determining levels of gene flow that influence the adaptability of species (Clobert, Baguette, Benton, & Bullock, 2012). Knowledge of dispersal can also to be vital to applied research that seeks to determine the conservation status of threatened species or the biosecurity risk of pest species, or for managing outbreaks of pests and disease vectors once invasions have taken place (Killeen, Knols, & Gu, 2003; Ouborg, Piquot, & Van Groenendael, 1999). Dispersal has traditionally been investigated by marking and tracking individuals across a landscape. However, these methods are often laborious and do not capture past dispersal patterns that produce gene flow. New methods and tools in landscape genetics can help understand past dispersal and gene flow patterns, including potential environmental parameters that limit dispersal (Schmidt, Filipović, Hoffmann, & Rašić, 2018; Watts et al., 2007). The advent of high-density sequencing technologies has also provided increased power for landscape genomic studies conducted at spatial scales fine enough to investigate dispersal discretely over generations, and to measure individual acts of movement (Schmidt, Rašić, et al., 2017).

Under Wright’s isolation by distance framework, the impact of dispersal on genetic variation in continuously distributed populations is described in terms of the neighbourhood size (*NS*), a product of parent-offspring dispersal variance and the ideal density of breeding adults within the dispersal area (Wright, 1946). Wright’s equation, *NS* = 4*πσ*^2^*d*, represents a key bridge between local demographic processes and broader patterns of genetic differentiation. As regards the ecology and genomics of a species, this bridge can be crossed in both directions, either using genomic differentiation to understand processes of dispersal and density, or alternately using estimates of density and dispersal to gain an insight into likely processes of genomic differentiation.

The link between *NS* and dispersal has been investigated using composite methodologies incorporating both individual genetic distances and mark-release-recapture (MRR) (Sahlsten, Thorngren, & Hoglund, 2008; Watts et al., 2007). However, for many species it is difficult to estimate density or dispersal using MRR methods. For instance, the small size of mosquitoes necessitates the use of field-sampled or laboratory-reared mosquitoes that are released for recapture from a central point (Honorio et al., 2003; Reiter, Amador, Anderson, & Clark, 1995). If marking materials or sampling protocols impact the fitness of released individuals, or if rearing conditions do not adequately represent release conditions, this methodology risks overestimating or underestimating dispersal distances. For example, *Aedes aegypti* (yellow fever mosquito) reared in suboptimal larval conditions may settle for ovipositing in suboptimal sites more readily than those reared in favourable larval conditions (Kaur, Lai, & Giger, 2003). Thus, if rearing conditions are unfavourable compared to conditions at the release site, released mosquitoes may disperse less than mosquitoes from the wild population under investigation.

MRR techniques can also be expanded to incorporate information from genetic inferences of kinship (Bravington, Skaug, & Anderson, 2016). This framework of close-kin mark-recapture (CKMR) uses a genetic ‘mark’ that extends beyond the individual by considering individuals to have genetically ‘marked’ their close kin, such that any kin sampled constitute an extended ‘recapture’ of the original individual. Kinship estimation can also be useful for investigating migration between populations, particularly in cases where migration rates have changed too recently to have produced corresponding changes in population genetic structure (Palsbøll, Zachariah, & Bérubé, 2010), such as might be expected in invasive species. While studies using microsatellites can often only confidently identify first-order kin relations (parent-offspring or full-sibling), the use of high density, genome-wide molecular markers can enable reasonably accurate assignment of individuals to second-order (e.g. half-sibling) and third-order (e.g. first cousin) groupings (Phillips, García-Magariños, Salas, Carracedo, & Lareu, 2012).

In this paper, we develop a methodology that uses genome-wide single nucleotide polymorphisms (SNPs) to estimate relatedness across three orders of kinship, and then uses the spatial distribution of these kin to estimate parent-offspring dispersal and related parameters. Our method inherits from CKMR, but exploits the fact that the spatial separation distance between each kin pair can be characterised as a set of distances representing past parental dispersal events (Figure 1), provided that the population is sampled at a specific point in time and from a set of sampling locations distributed continuously through space. By separating the composite life-histories associated with each kinship category into discrete parent-offspring dispersal events, we derive estimates of parent-offspring dispersal distances. Also, as most populations sampled continuously through space should have more second-order than first-order relations, and more third-order than second-order relations, this approach provides a means of investigating dispersal using a greater number of data points than by using first-order relatives alone, as dispersal distance estimates from first, second, and third-order relatives can all be incorporated into a single analysis. By using thousands of SNPs to estimate kinship and by integrating three orders of kinship into a single analysis, our methodology can be conducted with smaller sample sizes than those often required for kinship studies, a requirement which may have prevented more wide-spread adoption of kinship-based methodologies (Palsbøll et al., 2010).

We use the above methodology to investigate dispersal in *Ae. aegypti*, the primary vector of arboviral diseases such as dengue, chikungunya, and Zika (Morrison, Zielinski-Gutierrez, Scott, & Rosenberg, 2008). This highly anthropophilic mosquito is generally considered a weak disperser by flight (Harrington et al., 2005), though some MRR studies have reported flights over long distances (Honorio et al., 2003; Reiter et al., 1995). Given a local abundance of human hosts, the primary driver of dispersal in females is the search for suitable oviposition sites (Edman et al., 1998), and when undertaking ‘skip’ oviposition a female may oviposit at several sites during a single gonotrophic cycle (Reiter, 2007). Typically, females mate only once in their lives, while males may have multiple partners (Christophers, 1960). Despite its limited active dispersal, *Ae. aegypti* has managed to invade much of the global tropics over the past several centuries, dispersing passively along human trade and transport routes (Powell & Tabachnick, 2013).

**Figure 1.**
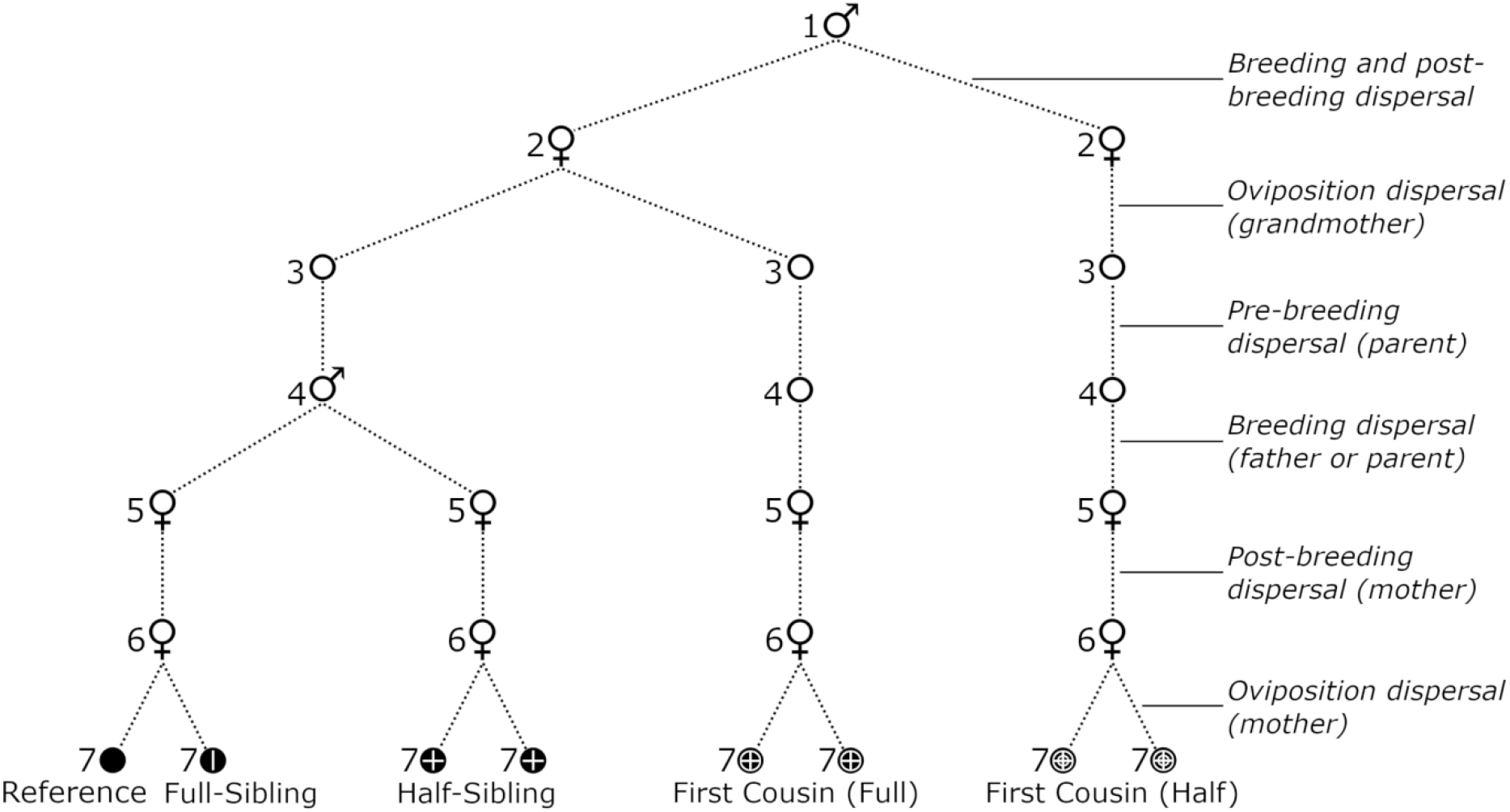
Past dispersal and breeding events underlying three orders of intragenerational kinship in *Ae. aegypti*. The set of past dispersal events for a given relatedness category is established by tracing from the reference along the dashed lines. Stages are as follows: (1) grandfather, (2) grandmother, (3) parent (as egg, larva, pupa), (4) parent (as adult, pre-mating), (5) mother (as adult, post-mating), (6) mother (ovipositing), and (7) sampled individual. Note that as females only mate once, all half-siblings are paternal half-siblings. Third order kin are assumed to be some mix of full-cousins and half-cousins.

We focus on dispersal across a residential site in Kuala Lumpur, Malaysia, as part of preparation for active interventions involving releases of *Ae. aegypti* transinfected with the bacterium *Wolbachia*. These interventions aim to replace uninfected *Ae. aegypti* with *Wolbachia*-infected mosquitoes that have a reduced potential to transmit dengue (Hoffmann et al., 2011; O’Neill et al., 2018; Schmidt, Barton, et al., 2017). The long-term persistence and spread of *Wolbachia* in *Ae. aegypti* will strongly depend on local *Ae. aegypti* dispersal characteristics (Hoffmann et al., 2014; Schmidt, Barton, et al., 2017; Turelli & Barton, 2017). Understanding intergenerational movement will also help inform strategies for responding to newly-detected *Ae. aegypti* incursions globally or to localised dengue outbreaks by chemical suppression of populations. However, applications of this methodology extend well beyond *Ae. aegypti* and other pest vectors. For instance, a key question in insecticide resistance management in agricultural pests concerns local movement of resistance alleles once resistance first arises, to help reduce resistance spread (Maino, Binns, & Umina, 2018). And in threatened species, understanding the spread of individuals across metapopulations can be vital in managing the species (Szczys, Oswald, & Arnold, 2017). Our approach to estimating dispersal and neighbourhood size therefore has numerous applications for both threatened and threatening species.

## Materials and Methods

### Mosquito sampling

*Aedes aegypti* were collected from Mentari Court in Petaling Jaya, Malaysia, between September 19^th^ and October 9^th^, 2017. Mentari Court consists of a set of 18-storey apartment blocks occupying a 500 × 250 m area, enclosing a vegetated area and a school (Figure 2). We deployed ovitraps on either the third or the fourth floor of each building, a sufficient height to avoid sampling *Ae. albopictus*, which are common at ground level in Mentari Court. We collected and stored *Ae. aegypti* larvae every week for three weeks. We divided the study region into nine sample sites across eight buildings (Figure 2) and sampled 18 mosquitoes from each site for DNA extraction, selecting six mosquitoes from different traps from each of the three weeks.

### DNA extraction and genotyping

DNA was extracted from the 162 *Ae. aegypti* using a Roche High Pure^™^ PCR Template Preparation Kit (Roche Molecular Systems, Inc., Pleasanton, CA, USA), with the addition of an RNase A digestion step. Extracted DNA was used to construct two RAD libraries containing 81 individuals each. We followed the double-digest Restriction-site Associated DNA sequencing protocol for *Ae. aegypti* developed by Rašić, Filipović, Weeks, and Hoffmann (2014), but selected a smaller size range of DNA fragments (350 – 450 bp) to accommodate the larger number of individuals in each library. The libraries were sequenced on the Illumina Hiseq3000 platform at the Monash Health Translation Facility, obtaining 100 bp paired-end reads and using a 20% Phi-X spike-in to reduce the impact of low diversity RADtags.

Following sequencing, barcoded reads were filtered, truncated to 80 bp, and demultiplexed using the process_radtags program in Stacks v2.0 (Catchen, Hohenlohe, Bassham, Amores, & Cresko, 2013). Paired and single-end reads were concatenated, and aligned to the *Ae. aegypti* nuclear genome assembly AaegL4 (Dudchenko et al., 2017) with Bowtie 2 (Langmead & Salzberg, 2012) using --very-sensitive alignment settings. We used the program ref_map to build a Stacks catalog and the program Populations to select SNP loci that were scored in at least 70% of mosquitoes. We filtered further with VCFtools (Danecek et al., 2011), retaining SNPs in Hardy-Weinberg equilibrium (P = 1e10^−5^) and with minor allele frequencies > 0.05, and thinned the remaining SNPs so that none was within 250 kbp of another. Thinning at this threshold in *Ae. aegypti* retains approximately 8 SNPs per map unit, a sampling density shown to largely eradicate linkage effects in SNPs (Cho & Dupuis, 2009). This set of 3,939 SNPs was used for all downstream analyses.

**Figure 2.**
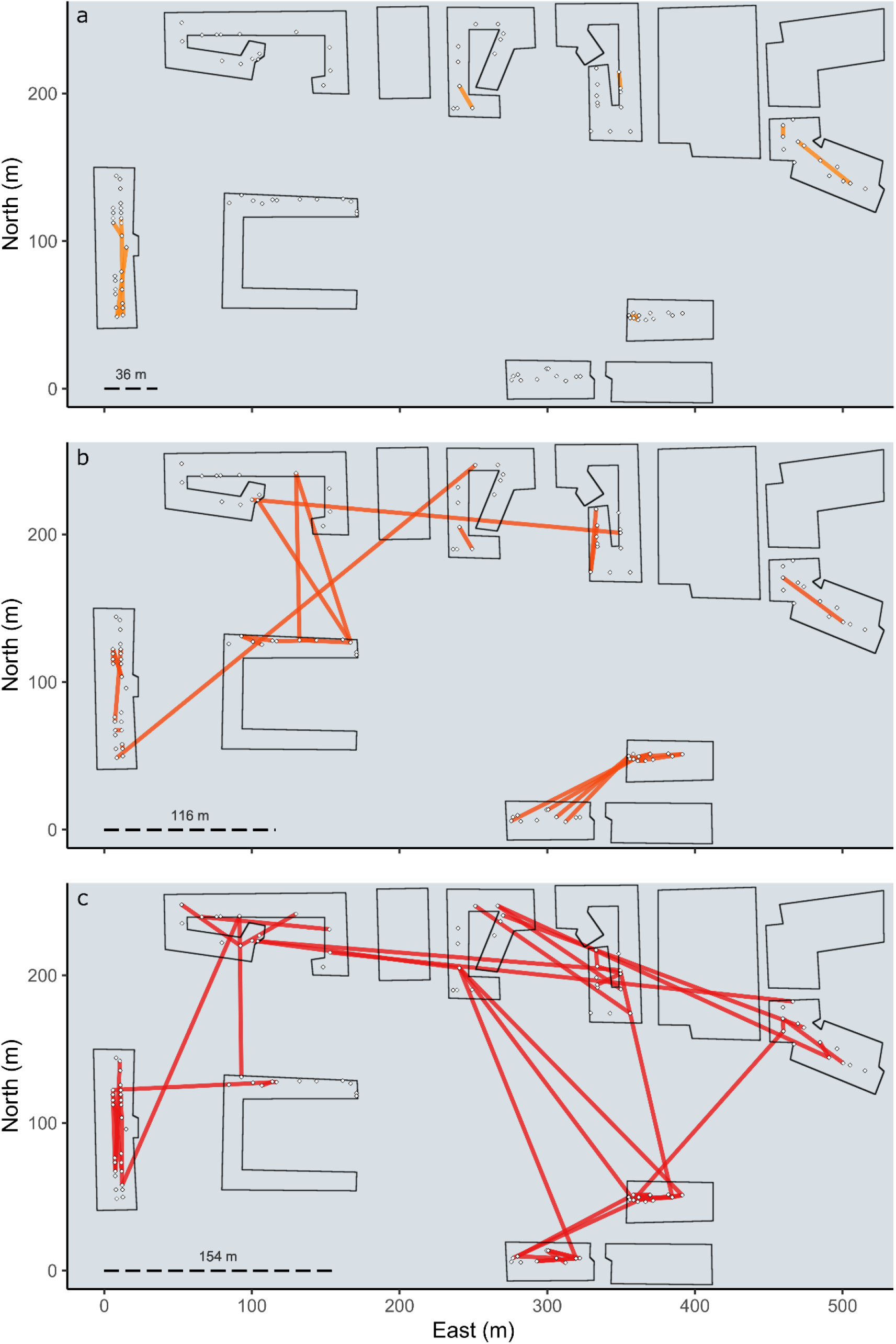
Mentari Court geography, trap placement, and kinship networks. Lines indicate pairs of full-siblings (a), half-siblings (b), and cousins (c). Rotated squares indicate ovitrap locations. Dashed lines show the radii within which 86.5% of pairs in that kinship category are expected to be found.

### Estimation of kinship categories and coefficients

To estimate the kinship category of each pair of mosquitoes, we calculated Loiselle’s kinship coefficient *k* (Loiselle, Sork, Nason, & Graham, 1995) in the program SPAGeDi (Hardy & Vekemans, 2002). Kinship coefficients represent the probability that any allele scored in both individuals is identical by descent (Wright, 1922), with theoretical mean *k* values for each kinship category as follows: full-siblings = 0.25, half-siblings = 0.125, full-cousins = 0.0625, half-cousins = 0.0313, second cousins = 0.0156, unrelated = 0. We ignored intergenerational kinship categories (e.g. uncle-niece), as the maximum of two weeks between the sampling of any pair of individuals would be insufficient time for the ontogeny of an additional generation (Christophers, 1960).

To assign pairs of individuals to relatedness categories across three orders of kinship (i.e. first cousins), we first used maximum-likelihood estimation in the program ML-Relate (Kalinowski, Wagner, & Taper, 2006) to identify first order (full-sibling) and second order (half-sibling) pairs. We used the *k* scores of pairs within the full-sibling and half-sibling datasets to calculate standard deviations for these categories.

However, ML-Relate is not configured to determine third order relationships (e.g. cousins). To determine cousins, we estimated a lower bound of *k* that separated first cousins from unrelated pairs and those of more distant kin groups. Here we define first cousins as including both full-cousins and half-cousins (Figure 1). We then produced simulated *k* scores for each kinship category, assuming that the *k* scores within each kinship category followed a normal distribution with a unique mean and standard deviation, and that these scores combined produced the entire distribution of *k* scores. Standard deviations for full-cousins, half-cousins, and second cousins were assumed to correspond to the standard deviation of the entire population after full-sibling and half-sibling pairs were removed.

Using the theoretical means and standard deviations of *k*, we randomly sampled 100,000 simulated *k* scores from each kinship category. In the initial pool of 13,041 empirical mosquito pairings, ML-Relate identified approximately 50 full-sibling and half-sibling pairs. Assuming that the data contained twice as many first cousin (full and half) pairings as sibling (full and half) pairings, and twice as many second cousin pairings as first cousin pairings, final sampling distributions were developed as follows: 100,000 unrelated, 2,000 second cousins, 500 full-cousins, 500 half-cousins, 250 half-siblings, and 250 full-siblings, giving a ratio of 400:8:2:2:1:1. This assumption is reasonable if the local population size is approximately constant; for a diploid population of constant size, an average of two offspring from any one individual will themselves have offspring. If we take *n* to be the number of larvae parented by one individual, the ratio of first cousin to sibling pairings would be approximately 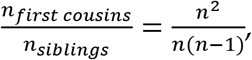 which ranges from 1.1 – 4 for 2 ≤ *n* ≤ 10. If we allow a slight population expansion with an average of 2.4 reproducing offspring, ratios from 1 ≤ *n* ≤ 10 produce values from 2.2 – 2.4.

To analyse how closely this distribution approximates the field data, we randomly sampled 10,000 simulated *k* scores from the above sampling distribution and plotted a histogram of this combined distribution and a histogram of the unrelated distribution against a histogram of 10,000 *k* scores from the empirical data. As the combined distribution matched the empirical distribution much more closely than the unrelated distribution (Supplementary Figure 1), we adopted it for kinship inference. Note that this unrelated distribution is not completely reflective of a natural population, as it assumes total random mating and zero population structure. It is a convenient null from which various models including this one can be developed and compared.

To determine a lower threshold of *k* to define a kinship category for first cousins, we plotted histograms of *k* scores for each kinship category, following the ratio described above (Supplementary Figure 2). We assigned *k* = 0.06 as the lower threshold to describe first cousin relationships. Individual pairs of *k* > 0.06 that were neither full-siblings nor half-siblings were much more likely to be first cousins (either full or half) than any other category.

### Inference of dispersal distributions

We treated separation distances between pairs of sampled kin as representative of multiple past dispersal events, following Figure 1. Full-sibling separation distances represent the dispersal of the mother during oviposition. Half-sibling separation distances represent the breeding dispersal of the father between matings, plus the post-mating dispersal of each mother between mating and oviposition, plus the ovipositional dispersal of each mother. Full-cousin separation distances represent the ovipositional dispersal of the grandmother, plus the pre-mating dispersal of each parent, plus the post-mating dispersal and ovipositional dispersal of each mother. Half-cousin separation distances are as those for full-cousins, plus the breeding dispersal of the father between matings and the post-mating dispersal of each mother between mating and oviposition. In all cases, we considered dispersal to operate across a two-dimensional plane, and ignored any differences in sampling altitude between buildings.

Axial standard deviations correspond in form to the dispersal component of *NS* (Wright, 1946). We calculated axial standard deviations from the distributions of separation distances for each kinship category. To do this, we (i) projected distances onto a polar coordinate system with a random angle of rotation, (ii) converted the distances to one-dimensional vectors by multiplying each distance by the cosine of its rotation angle and (iii) calculated the standard deviation of the resulting distribution. The final axial standard deviations were estimated by applying steps i) – iii) to each kinship category. See Supplementary Text for details.

### Determination of parent-offspring dispersal

Parent-offspring dispersal is described by Wright (1946) as the distribution of parents at some phase of the life cycle relative to offspring at the same phase. These events serve to produce geographical distributions of not only full-siblings and half-siblings but full-cousins and half-cousins as well (Figure 1). Specifically, the full-cousin and half-cousin dispersal distributions are produced by combining the parent-offspring dispersal distributions with the full-sibling and half-sibling distributions respectively. As the variance of the normal distribution formed by the combination of two normal distributions is equal to the sum of the component variances, we can infer parent-offspring axial dispersal, σ_PO_, with the following equations:

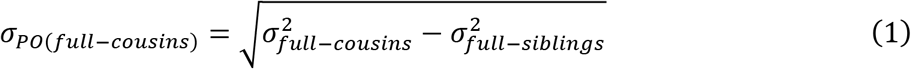

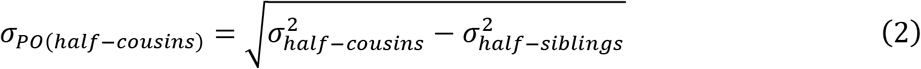

As we assume a mixed distribution of full-cousins and half-cousins in the kinship pairings, these represent the upper (full-cousin) and lower (half-cousin) bounds of parent-offspring dispersal estimates using the first cousin distribution. Assuming a 50:50 ratio of full-cousins and half-cousins in our data, the mean parent-offspring axial dispersal can be approximated by the following equation:

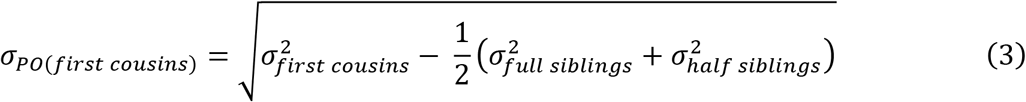

We use equation (3) for all future calculations of parent-offspring dispersal. For each axial dispersal distribution (σ), 2σ represents the effective radius of dispersal, within which 86.5% of dispersed individuals are expected to be found (Wright, 1946). In the case of parent-offspring dispersal, this constitutes the radius of a circle defining Wright’s neighbourhood area, the spatial component of the *NS* estimate within an isolation by distance framework.

### Geographical genetic structure and neighbourhood size

We performed distance-based redundancy analyses (dbRDA) to quantify the effects of geographical distance, sampling week, and building residency on patterns of genetic distance-observed among *Ae. aegypti* at Mentari Court. For these tests, we resampled the dataset so that no full-siblings or half-siblings were included, retaining 130 individuals. This was necessary as the inclusion of sibling pairs can bias estimates of population structure (Goldberg & Waits, 2010). We were interested in whether genetic distances were spatially structured at this scale, and if so whether structure best fit a pattern of isolation by distance, isolation by time, isolation by building residency, or some combination of the three. All analyses were performed with the R package “vegan” (Oksanen et al., 2010), and all geographical distances were transformed with a natural logarithmic function.

We used dbRDA to build three models. The first (“dist”) tested for effects of only geographical distance on genetic distance. The other models tested for effects of geographical distance on genetic distance after conditioning for the effects of sampling week and building (“dist | *env*”), and for effects of sampling week and building on genetic distance after conditioning for the effects of geographical distance (“env | *dist”*). We quantified the marginal explanative power of each variable in each of the dbRDAs using ANOVA with 10,000 permutations. The functions *capscale* and *anova.cca* were used for dbRDA and ANOVA respectively.

As dbRDA requires independent variables to have one-dimensional inputs, we first transformed the matrix of pairwise geographical distances into eight principal components (PCs) using the function *pcnm*. Sampling time was modelled as a continuous variable of integers corresponding to the first, second, and third week of sampling, while building residency was modelled categorically. We used a pairwise matrix of Rousset’s *a* scores (Rousset, 2000) as the dependent variable, calculated in SPAGeDi. For geographical distance in the dbRDAs “dist | *env”* and “env | *dist”*, we used only the PCs that were significant in the dbRDA “dist” (P < 0.006 following Bonferroni correction).

In a population experiencing isolation by distance, *NS* is estimated as the inverse of the slope of the regression of pairwise genetic distance against the natural logarithm of geographical distance (Rousset, 2000). That is, *NS* = *b*^−1^, where *Genetic distance* = *a* + ln(*Geographical distance*) * *b*. We used the R function “lm” to perform three of these regressions. The first, using all non-sibling pairs of individuals, we used to estimate *NS* following the above. For the other two regressions we considered only within-building pairings and between-building pairings, to investigate isolation by distance within and between buildings respectively.

## Results

### Kinship inference

Using the 3,939 SNPs retained after filtering, we identified 13 full-sibling pairs and 34 half-sibling pairs using ML-Relate. We also designated 51 pairs with *k* > 0.06 as first cousins. Mean separation distance for full-siblings was 18.1 m, for half-siblings it was 48.6 m, and for cousins it was 75.1 m (Table 1). Supplementary Figure 3 shows the spatial distribution of *k* scores across geographical distance for all kinship categories as scatterplots (3A) and density histograms (3B).

**Table 1.**
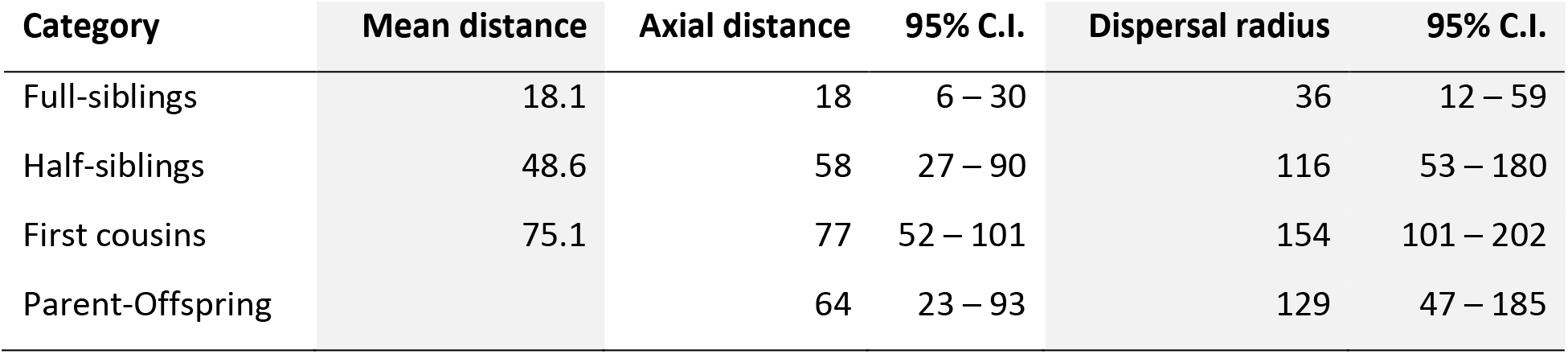
Estimates of dispersal distances in metres for each kinship category. Mean distance describes empirical distances. Axial distance describes the axial standard deviation of a bivariate normal distribution of dispersal distance. Dispersal radius describes the radius within which 86.5% of dispersed individuals are expected to be found.

### Determination of parent-offspring dispersal via kinship distributions

For each kinship category, we used geographical distances of separation between kin to construct a bivariate normal distribution, from which we calculated the axial standard deviation of dispersal (Table 1). Using the full-sibling, half-sibling, and first cousin standard deviations, we estimated parent-offspring axial dispersal as 64 m (23 – 93 m; 95% C.I.). The radius of effective parent-offspring dispersal is equal to double the parent-offspring axial standard deviation, coming to 129 m (47 – 185 m; 95% C.I.). This corresponds to neighbourhood area, the circle within which 86.5% of parent-offspring dispersal occurs.

Following the process for parent-offspring dispersal, we generated dispersal radii for each kinship distribution. Referencing the axial standard deviations in Table 1, we derived a dispersal radius of 36 m (12 – 59 m; 95% C.I.) for full-siblings, representing ovipositional dispersal; a dispersal radius of 116 m (53 – 180 m; 95% C.I.) for half-siblings, representing breeding and ovipositional dispersal; and a dispersal radius of 154 m (104 – 202 m; 95% C.I.) for first cousins (see Figure 1). Additionally, through simulations of the parent-offspring axial standard deviation, we estimated the median parent-offspring dispersal distance as 75 m, which represents the range within which 50% of dispersal events lie.

Pairwise locations of all full-siblings, half-siblings, and first cousins are shown in Figure 2. All full-sibling pairs were found in the same building (Figure 2a). Half-sibling pairs were mostly found in the same building or adjacent buildings, although two pairs were found between nonadjacent blocks (Figure 2b). First cousin pairs were also mostly found in the same building or adjacent buildings, but were distributed between nonadjacent buildings more often than half-siblings (Figure 2c).

### Geographical genetic structure and neighbourhood size

The dbRDA evaluating the effects of geographical distance on genetic structure (“dist”) indicated that 3 of 8 PCs were within the significance threshold (Bonferroni-corrected critical value: P < 0.006). These PCs were used as the geographical distance covariate in the dbRDAs env | *dist* and dist | *env* (Table 2). After conditioning for the effects of geographical distance (Bonferroni-corrected critical value: P < 0.025), env | *dist* showed that building residency had a significant effect on patterns of genetic structure (P = 0.002; F = 1.67; 7 d.f.), while sampling week did not (P = 0.854; F = 0.690; 2 d.f.). After conditioning for the effects of building residency and sampling week, the 3 PCs in dist | *env* showed no significant association between geographical distance and genetic distance (P = 0.0901; F = 1.155; 3 d.f.). As building residency and geographical distance were necessarily correlated, these results indicate a significant effect of building residency independent of geographical distance, and an uncertain effect of geographical distance.

**Table 2.**
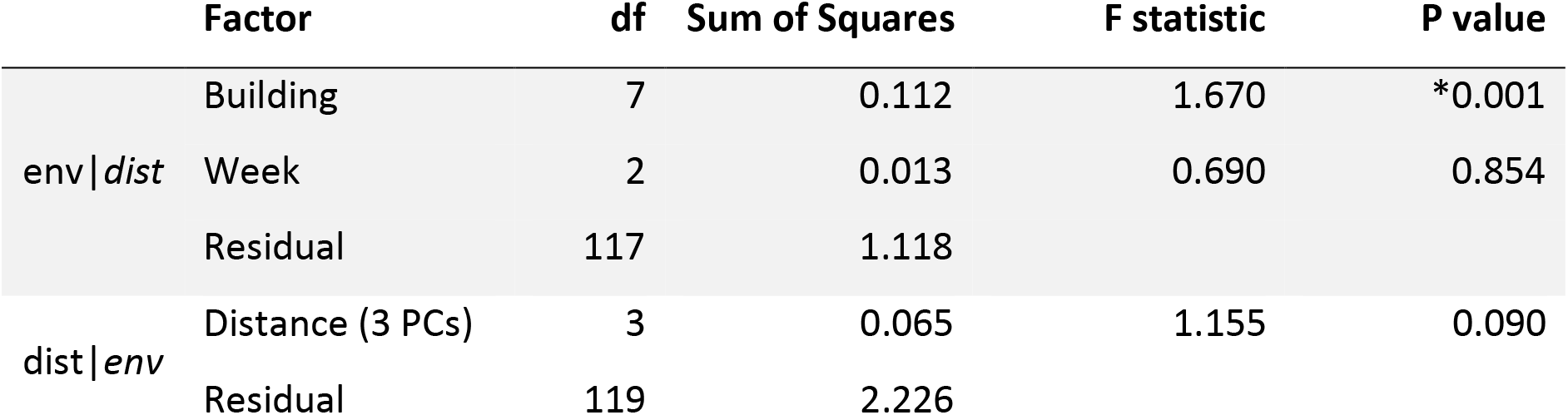
Results of dbRDAs. These test effects of building residency and sampling week on genetic distance after conditioning for geographical distance (env | *dist*), and effects of geographical distance after conditioning for building residency and sampling week (dist | *env*). Asterisk denotes significance at the Bonferroni-corrected critical value of P < 0.025 (env | *dist*) or P < 0.05 (dist | *env*).

We explored the effects of geographical distance further using linear regressions. The three linear regressions calculated for all non-sibling pairs (P < 2e^−16^; R^2^ = 0.0090; slope = 0.0037), all non-sibling pairs within the same building (P = 0.012; R^2^ = 0.0062; slope = 0.0041), and all non-sibling pairs in different buildings (P = 1.3e^−09^; R^2^ = 0.0048; slope = 0.00484), all showed significant positive associations between geographical and genetic distances (Supplementary Figure 4). These isolation by distance patterns indicate that *Ae. aegypti* populations are spatially structured within buildings as well as between buildings, show slightly stronger structuring between buildings than within buildings, and show a clear effect of geographical distance independent of building residency. Considering the size of buildings at Mentari Court, this implies isolation by distance at spatial scales of less than 200 m. Using the slope of the regression for all non-sibling pairs, we calculated *NS* as 268 *Ae. aegypti* (222 – 345; 95% C.I.). This constitutes the effective number of breeding individuals inhabiting a circle of radius 258 m, which is two times the axial standard deviation of parent-offspring dispersal (Wright, 1946).

## Discussion

This study has developed a methodology that uses genomic inferences of kinship to estimate parent-offspring dispersal and related population parameters, and applied this methodology to investigate an urban population of the invasive disease vector *Ae. aegypti*. From 162 sequenced individuals we identified 98 pairs of kin across three orders of kinship. By limiting the duration of sampling and by sampling only eggs, we were able to ignore intergenerational kin categories, and thus assigned the 98 pairs to intragenerational categories: full-sibling, half-sibling, full-cousin, and half-cousin. After transforming the spatial distances separating kin pairs within each category into axial standard deviations, we isolated the parent-offspring component of dispersal, which corresponds to the difference between the axial standard deviations of full-siblings and full-cousins and of half-siblings and half-cousins. Using our final estimate of parent-offspring dispersal, we calculated neighbourhood area (129 m), median parent-offspring dispersal distance (75 m), and radius of ovipositional dispersal within a gonotrophic cycle (36 m). Our additional analyses with dbRDA and linear regression revealed isolation by distance both within and between buildings, and provided an estimate of *NS* (Wright, 1946) of 268 individuals. These results indicate the scale of intergenerational dispersal and population structuring in *Ae. aegypti*, and thus also the scale at which efforts must be focused to restrict new invasions, suppress or transform established populations, and respond strategically to arboviral outbreaks. With appropriate modifications where required, our methodology will be readily applicable to studies of other species of interest such as pests and those of conservation significance.

Our estimates of parent-offspring dispersal, combined with the results of the dbRDAs and linear regressions, suggest that the *Ae. aegypti* population of Mentari Court is hierarchically structured. The linear regressions indicate isolation by distance patterns both within (< 200 m) and between buildings, while the dbRDAs showed that building residency had an effect on structure even after accounting for geographical distance. These results suggest dispersal within buildings is less restricted than dispersal between buildings, and that dispersal between adjacent buildings is more common than between nonadjacent buildings. This interpretation accords with kin observations, wherein most kin pairs found in nonadjacent buildings were cousins (Figure 2), which are a result of multiple parent-offspring dispersal events (Figure 1). This also accords with our estimate that 86.5% of parent-offspring dispersal occurs within a circle of radius 129 m, which corresponds in scale to movement within buildings and between adjacent buildings but not between nonadjacent buildings. Thus, dispersal of *Ae. aegypti* between nonadjacent buildings at Mentari Court is likely to be a multigenerational process. This equates to a matter of months for *Ae. aegypti*, which typically has 10 – 12 generations per year (Christophers, 1960).

These findings are broadly consistent with previous estimates from MRR studies which describe a tendency of *Ae. aegypti* to stay within the building of its release, and when dispersing farther rarely travel more than 150 m (Harrington et al., 2005; Maciel-de-Freitas, Codeco, & Lourenco-de-Oliveira, 2007; Ordonez-Gonzalez, Mercado-Hernandez, Flores-Suarez, & Fernandez-Salas, 2001; Russell, Webb, Williams, & Ritchie, 2005). Occasional long-distance dispersal > 500 m has also been recorded through MRR (Honorio et al., 2003; Reiter et al., 1995); while these dispersal distances are larger than the sampling scale of this study, they have been observed in genomic studies (Schmidt et al., 2018). Likewise, our finding that buildings or the space between them acts as a dispersal barrier is consistent with observations of fine-scale dispersal barriers in *Ae. aegypti* (Hemme, Thomas, Chadee, & Severson, 2010; Schmidt et al., 2018). The 18-storey high-rise blocks of Mentari Court are populated with *Ae. aegypti* up to the top floor, and are likely to provide adequate within-building opportunities for mating, human blood-feeding, and oviposition, with dispersal within buildings likely to be safer and easier than crossing the more ‘hostile’ open spaces between buildings. Future work could examine dispersal patterns in building complexes of fewer storeys to test if these lead to higher rates of movement between buildings, which could result from smaller buildings having fewer oviposition sites. Future work could also investigate vertical dispersal between floors, which was not considered in this study.

If movement patterns of *Ae. aegypti* through Mentari Court are representative of high-rise building complexes generally, we expect these findings to have implications for understanding *Ae. aegypti* as a disease vector, as an invasive species, and as a target of biological control initiatives. In the event of an arbovirus outbreak, the initial spread of the virus by female *Ae. aegypti* will likely be restricted to within buildings or between nearby buildings. Spread by females to more distant buildings would likely take multiple generations, or multiple gonotrophic cycles of a single female, and thus proceed more slowly. Following a new detection of invasive *Ae. aegypti*, we expect limited active dispersal from the point of introduction, though passive dispersal by human transport may spread the incursion more quickly. Finally, considering releases of *Wolbachia*-infected *Ae. aegypti* at Mentari Court, we expect that successful invasion of one building with *Wolbachia* is unlikely to result in the invasion of other buildings, as movement between buildings may be insufficiently common for the invasion to surpass a critical threshold frequency of 0.25 – 0.35 (Schmidt, Barton, et al., 2017; Turelli & Barton, 2017). However, limited dispersal between buildings is also likely to ensure that a successfully invaded building remains so, as few infected *Ae. aegypti* will leave and few uninfected *Ae. aegypti* will arrive.

Using the regression between individual genetic distance and the natural logarithm of geographical distance (Rousset, 2000), we estimated *Ae. aegypti NS* at 268 individuals. Combining this with our parameter for parent-offspring dispersal (σ) of 129 m, we can estimate the ideal density *d* of *Ae. aegypti* breeding adults as 1281 km^−2^, using Wright’s equation for *NS* under isolation by distance, *NS* = 4*πσ*^2^*d* (Wright, 1946). In interpreting these values, it is essential to be clear about the distinction between *NS* and effective population size (*N_e_*). *NS* is a parameter representing breeding structure in an isolation by distance framework, and relates directly to dispersal, while *Ne* is dependent on local species abundance and thus habitat availability (Nunney, 2016). These parameters are therefore expected to estimate different numbers of individuals. An extensive study of *Ae. aegypti* estimated average *N_e_* of 400 – 600 across different sites and timepoints (Saarman et al., 2017), while census size estimates range from 900 – 5,500 individuals (Carvalho et al., 2015; Lounibos, 2003; Sheppard, Macdonald, Tonn, & Grab, 1969). Relative census sizes of populations can be estimated using variations in *N_e_* and may improve our ability to understand *Ae. aegypti* population characteristics. Census size estimates are often five to ten-fold larger than those of *N_e_* (Luikart, Ryman, Tallmon, Schwartz, & Allendorf, 2010; Palstra & Fraser, 2012); were a similar ratio to hold for *NS*, we would expect to find between 1,340 and 2,680 *Ae. aegypti* within the dispersal radius, a figure consistent with typical adult census population sizes for this species. All of this emphasizes how combining kinship-based dispersal estimates with more widely-used, distance-based investigations of genetic structure can improve our understanding of local population and breeding processes in *Ae. aegypti*.

Apart from the introduction of novel oviposition sites through ovitraps, our methodology of inferring dispersal through genetic relatedness leaves dispersal processes essentially undisturbed, an advantage over MRR studies. Accordingly, genomic inferences of dispersal may more accurately reflect the true biological processes occurring within the study site. An additional advantage of kinship-based methodologies is their capacity to investigate population processes such as dispersal among populations in which migration rates have changed too recently to have produced corresponding changes in population genetic structure (Palsbøll et al., 2010), such as might be expected in invasive species and those with habitats impacted by humans. Ongoing decreases in sequencing costs means that the approach is becoming feasible for a wider range of species, particularly as our methodology parameterises dispersal using estimates from three orders of kinship, and does not depend on large sample sizes required for many kinship-based methodologies. When stringent requirements on sample size are relaxed, more widespread adoption of kinship-based methodologies seems likely (Palsbøll et al., 2010), particularly since current genetic markers allow for more accurate estimation of kinship categories than previously-used markers (Hauser, Baird, Hilborn, Seeb, & Seeb, 2011). In this study we sequenced 162 *Ae. aegypti* on only two sequencing lanes, which provided sufficient read depth and breadth to identify 98 pairs of close kin.

Our methodology is applicable to a range of species, including those of medical, economic, or conservation significance. However, differences in breeding and dispersal ecologies between species will alter the way that the method is applied. For instance, for species where females mate more than once, additional steps will be required to separate half-sibling dispersal from maternal dispersal. Similarly, species with sex-biased dispersal will require additional steps when decomposing axial standard deviations into components. Also, while our method is well-suited to studying rare species of conservation significance, in which direct population manipulation through MRR may not be possible, these studies will likely have low sample size and involve inbred individuals (Hedrick & Kalinowski, 2000). These conditions may make it more difficult to estimate kinship using genomics, though we note that parts of our methodology can equally be applied using kinship estimates derived through other means such as visual tracking of relatives.

The kinship-based inference processes described here have close affinities with the work of Bravington et al. (2016), and constitute an important extension to CKMR. Using genome-wide SNPs, we have been able to extend the CKMR methodology to cover three orders of kinship, estimating relatedness to the level of half-cousin. Likewise, this study has developed an approach for estimating intergenerational dispersal through the genomic marking of unsampled parents and grandparents. This will be particularly helpful for studies of species with traits such as small size or high population density that make them unsuitable for standard MRR experiments,. For example, monitoring of rare mammals is often carried out by the scats they leave, with recent work using DNA from scats to infer *N_e_* from kinship (Skrbinšek et al., 2012). As methods for retrieving DNA from scats improve (Schultz, Cristescu, Littleford-Colquhoun, Jaccoud, & Frère, 2018), investigating kin networks and intergenerational dispersal with high density markers may become increasingly viable for achieving conservation objectives.

## Conclusions

We developed and applied a methodology to investigate dispersal and breeding dynamics in *Ae. aegypti* from the spatial distribution of 98 pairs of kin across three orders of kinship. We also observed genetic structure within and between buildings at Mentari Court, with most movement between nonadjacent buildings taking place over multiple generations. These findings are being considered in the design of future *Wolbachia* releases at Mentari Court and other high-density urban sites, and they will also inform protocols for the effective deployment of resources in response to disease outbreaks. Our methodological approach for investigating dispersal and breeding dynamics will be readily adaptable to future studies of dispersal in a range of species.

## Acknowledgements

This work was funded by the National Health and Medical Research Council (Program Grant no. 1132412; Fellowship Grant no. 1118640), and the Wellcome Trust (Grant no. 108508). We thank three anonymous reviewers for their comments.

## Data Accessibility

Sorted .bam files and geographical locations for 162 *Aedes aegypti* have been archived at the Sequence Read Archive at NCBI Genbank, with accession number PRJNA542421.

## Author Contributions

Moshe Jasper: Performed research, wrote the paper, analysed data, contributed new analytical tools

Thomas L. Schmidt: Performed research, wrote the paper, provided supervision

Nazni W. Ahmad: Designed research, performed research

Steven P. Sinkins: Designed research, provided funding

Ary A Hoffmann: Designed research, provided funding, provided supervision

## Supplemental Figures for

**Supplementary Figure 1.**
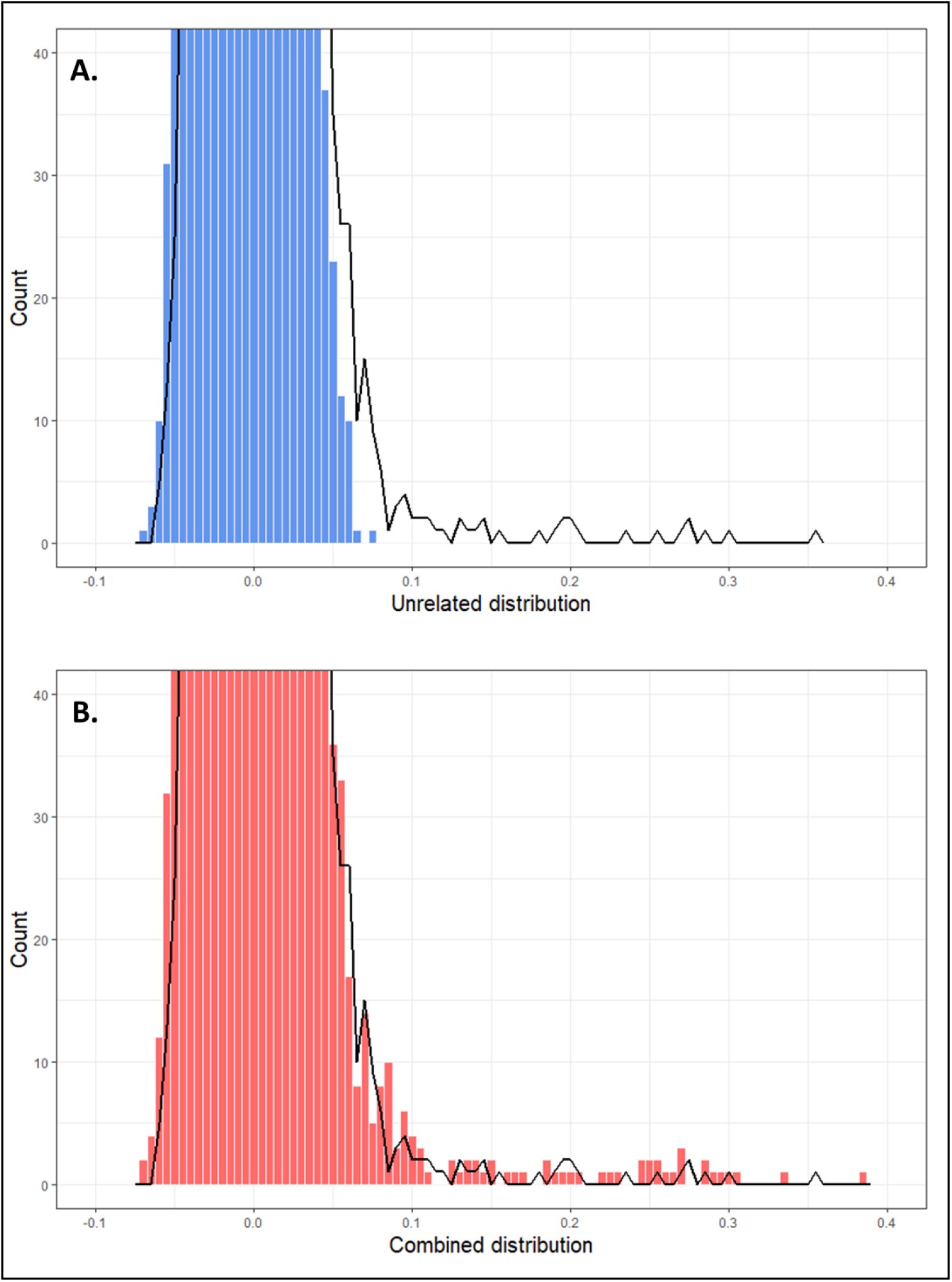
**A** Histogram of unrelated sampling distribution (blue) compared to that of field data (black line). **B** Histogram of combined sampling distribution (red) compared to that of field data (black line). The combined sampling distribution approximates the field data much more closely than does the unrelated.

**Supplementary Figure 2.**
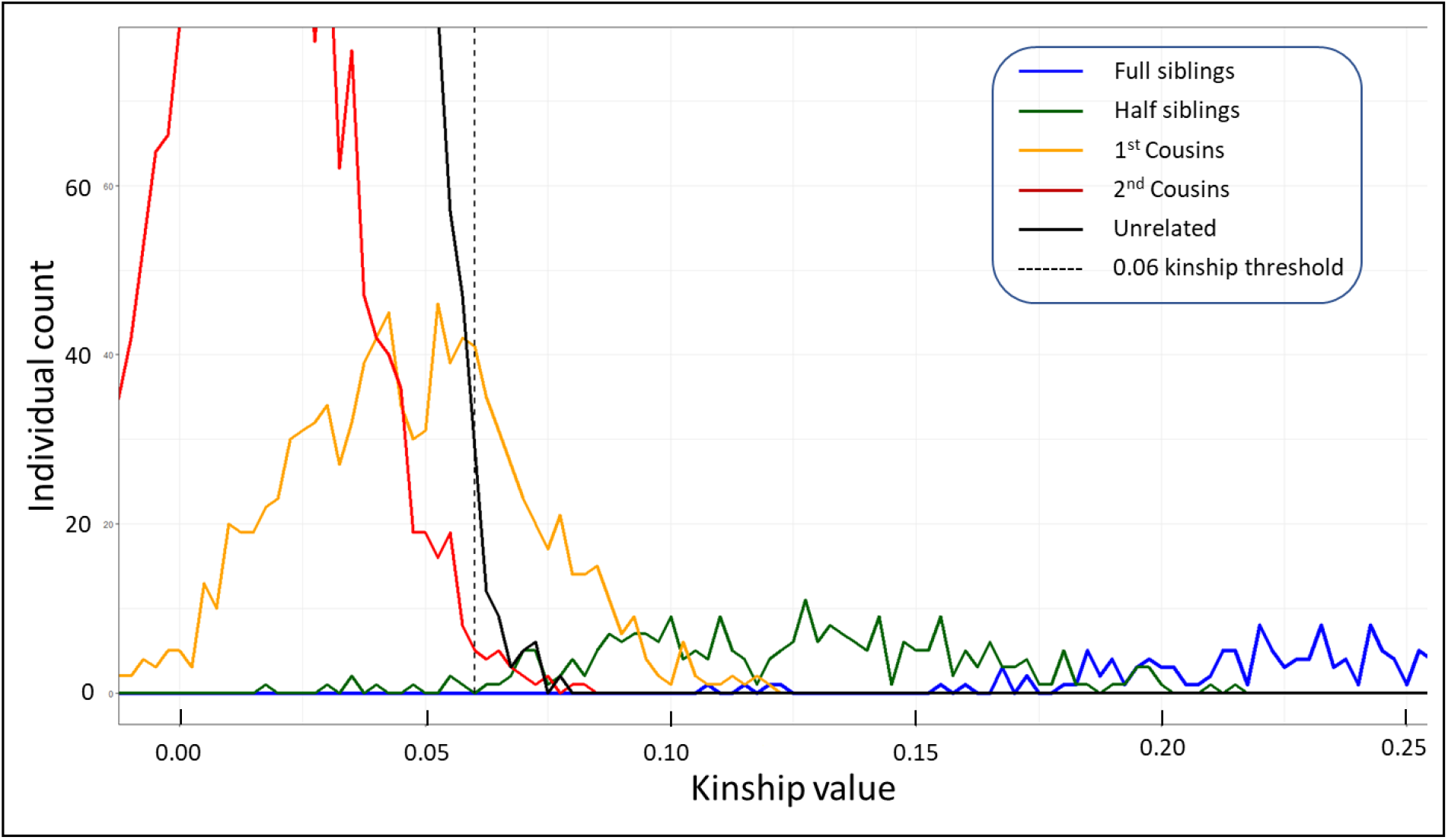
Frequency polygons showing distributions of simulated kinship (*k*) scores. Colours indicate unrelated (black), second cousins (red), first cousins (orange), half-siblings (green), and full-siblings (blue). Dotted vertical line shows the *k* = 0.06 lower bound for first cousins. Based on these distributions, we no longer considered second cousins.

**Supplementary Figure 3.**
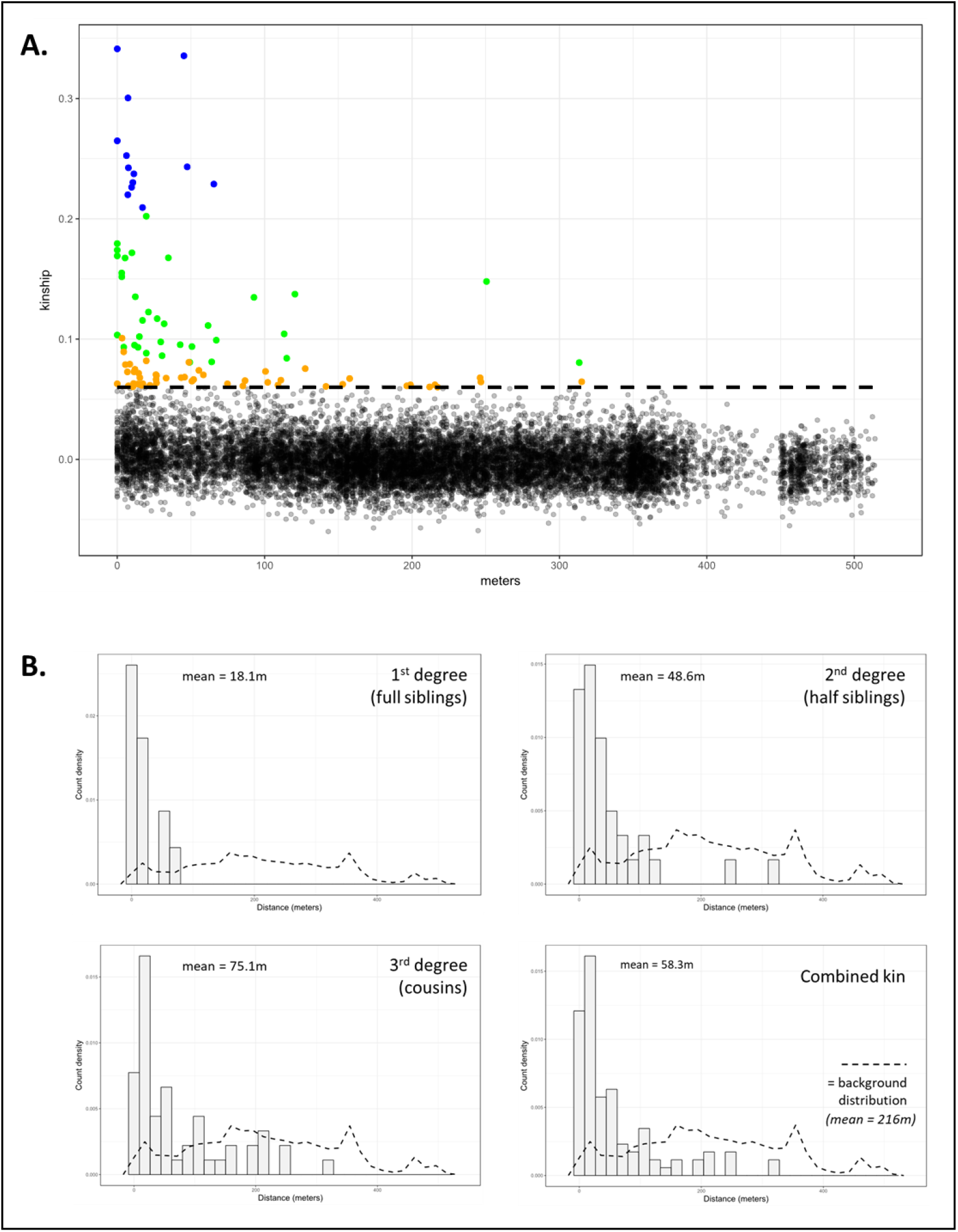
Kin assignment and distribution across distance. **A** Pairwise Loiselle’s *k* scores between all individuals relative to distance of separation. Blue = full-siblings, green = half-siblings, orange = cousins, black = unrelated. Dotted line at *k* = 0.06 shows the lower bound for first cousins. **B** Histograms of density of each order of kinship (and combined) relative to distance (m). The density distribution of all possible pairwise combinations is shown as a dotted line in each panel for reference.

**Supplementary Figure 4.**
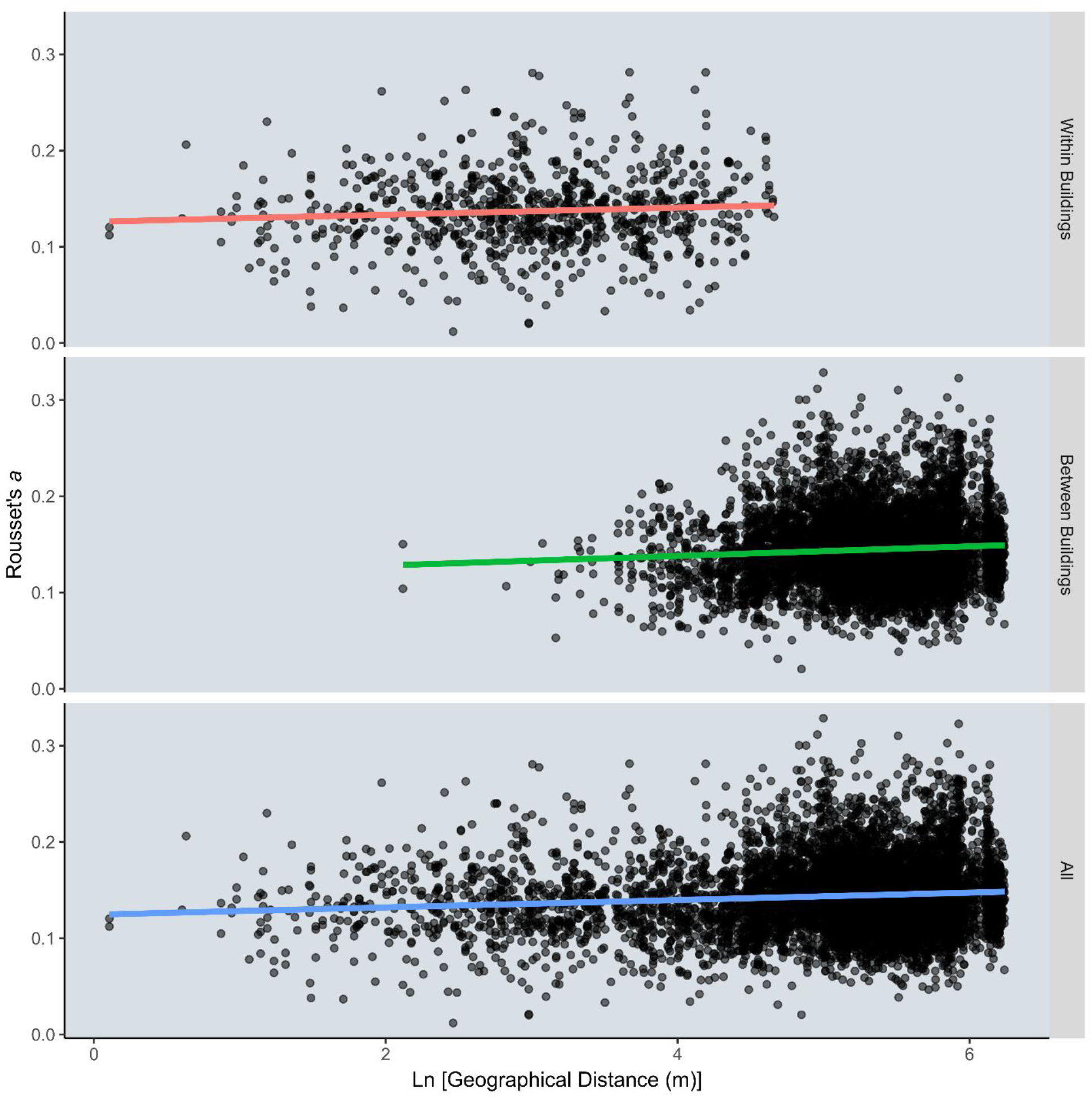
Regressions of pairwise genetic distance (Rousset’s a) and the natural logarithm of geographical distance for all non-sibling pairs. Regressions for all non-sibling pairs (P < 2e^−16^; R^2^ = 0.0090; slope = 0.0037), all non-sibling pairs within the same building (P = 0.012; R^2^ = 0.0062; slope = 0.0041), and all non-sibling pairs in different buildings (P = 1.3e^−09^; R^2^ = 0.0048; slope = 0.00484) all showed significant positive associations between geographical and genetic distances.

## Calculation of Axial Standard Deviations of Dispersal

### Theory

To enable the calculation of axial standard deviations, each dispersal process is modelled as a biva-riate normal distribution with a mean of (0, 0) and rotational symmetry, allowing both axial standard deviations to be equivalent. Under these assumptions, the following procedure derives axial standard deviations from the distributions of separation distances for each kinship category:

1. For each separation distance between kin, assign a random angle of rotation. Applied aggregatively, this removes any directional sampling biases in the data while projecting the distances onto a polar coordinate system.
2. Convert the distances to one-dimensional vectors by multiplying each distance by the cosine of its rotation angle. For a polar coordinate system, this flattens the available distance information to a one-dimensional distribution centred around zero – an axial distribution.
3. Calculate the standard deviation of the resulting distribution.
4. For statistical inference by bootstrapping, apply the above steps to 1,000 subsamples produced by resampling with replacement from within the kinship category, with angles randomised in each subsample. The average of the resulting 1,000 standard deviations is an estimate of the axial standard deviation, while the 2.5% and 97.5% quantiles of the distribution define 95% confidence intervals for the estimate.
5. For Parent-Offspring inference, use equation 3 (main text) to derive an estimate from axial standard deviations of the other categories, bootstrapping on final P-O estimates.

### Code

#### Dependencies

The code to calculate axial deviations depends on the dplyr package https://cran.r-project.org/web/packages/dplyr/index.html

#### Data setup

To run the following code, pairwise kinship and distance data must be set up as a dataframe, with each row corresponding to a separate pairwise comparison between two individuals.

A variety of fields could prove useful in this frame, but typically will at least include Loiselle’s kinship (*k*), genetic distance, and geographical distance (in this example, the $meters field).

During kinship filtering, this main dataframe has been split into several smaller kinship groups (dataframes) labelled fullsibs, halfsibs, and cousins. Each dataframe contains all pairwise comparisons classified under that kinship grouping.

#### Axial Distance Functions

The first function, **axialdist**, takes a distribution of distances and calculates an axial standard deviation with random angles from it (no replicates, unless specified). This corresponds to steps 1 – 3 in the paper.

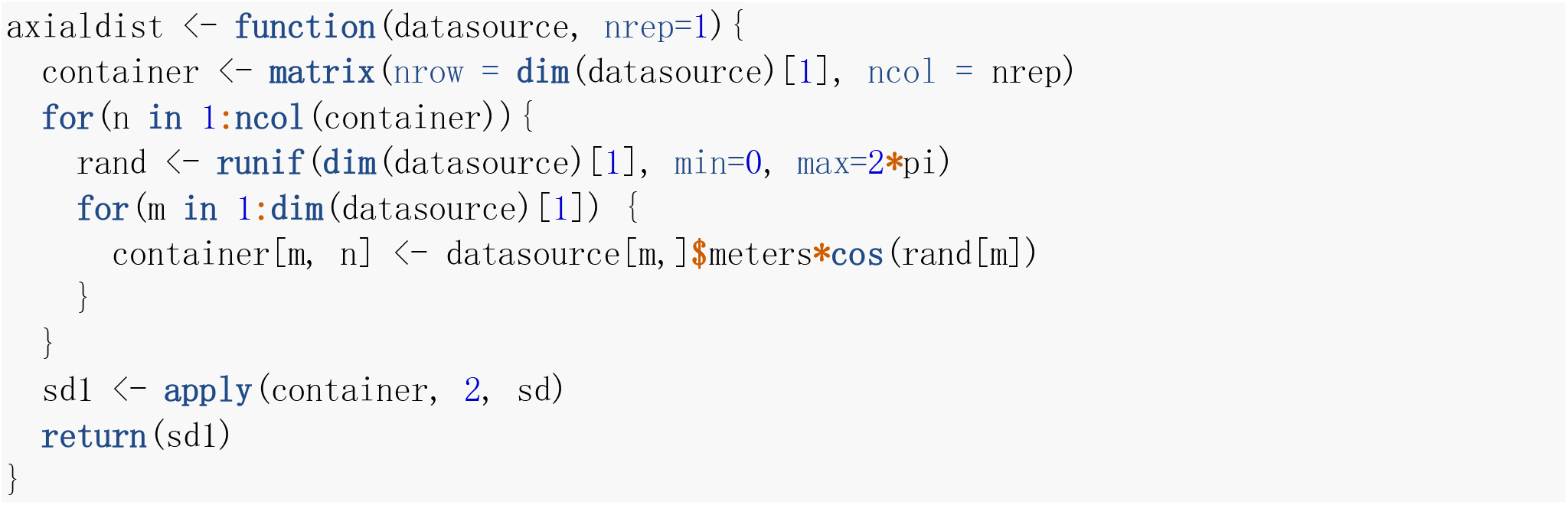

The second function, **quantax**, applies the first to 1,000 samplings of a kinship group and returns the quantiles, including those required for 95% confidence interval.

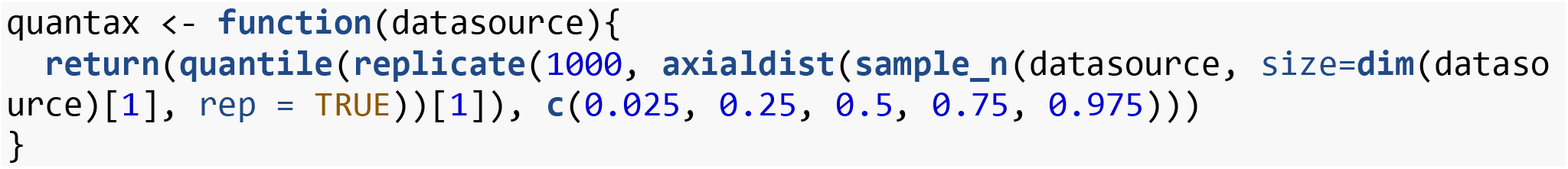

The third function, **po_quantile**, returns quantiles for parent-offspring axial standard deviation, including those required for 95% confidence intervals, through resampling the underlying distributions 1,000 times. Note that as sampling distributions for cousins overlap with those of halfsibs, a small number of trials result in Zero Division Errors, which can be discarded.

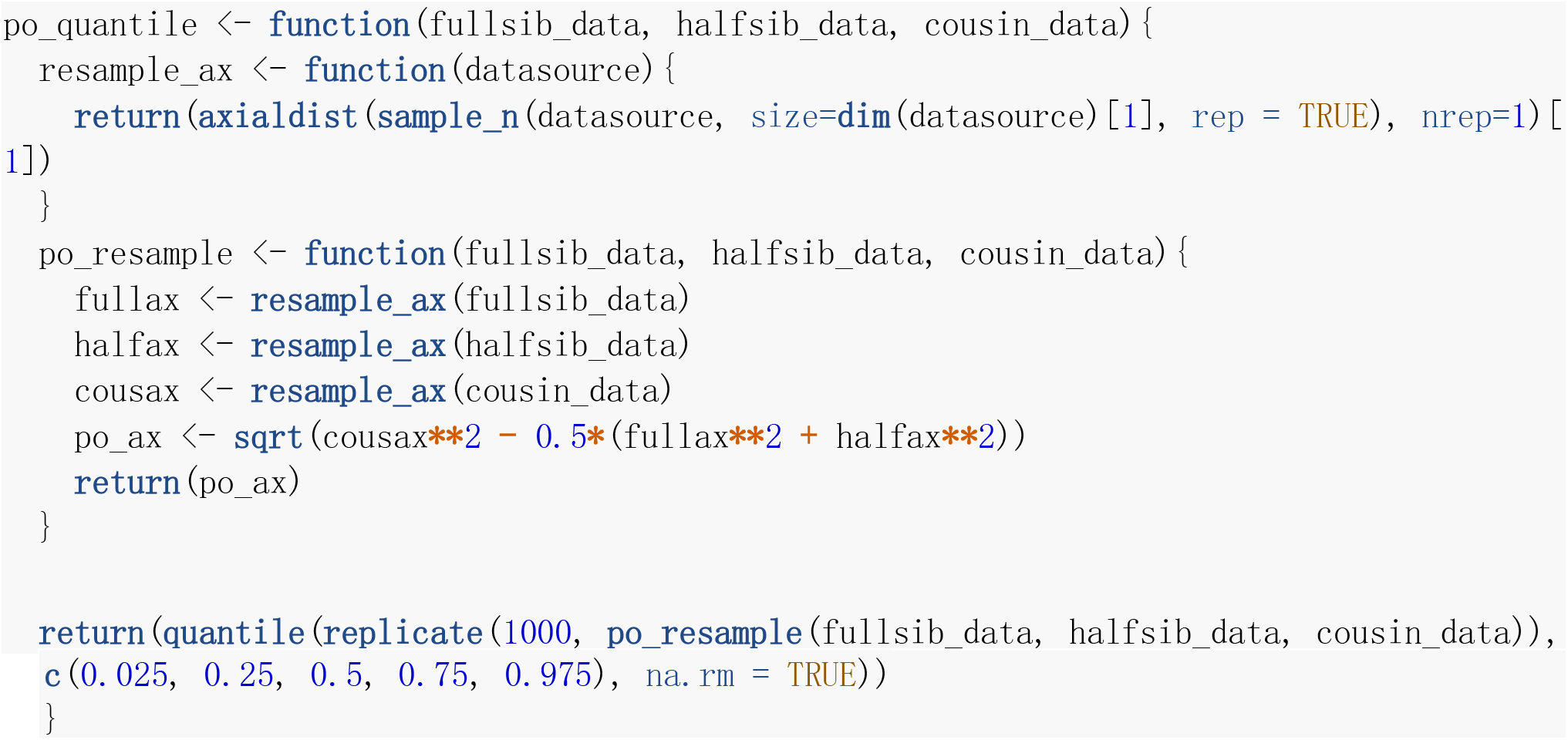

#### Final code

Using the above data structures and functions, final axial dispersal parameters are **calculated as shown below.**

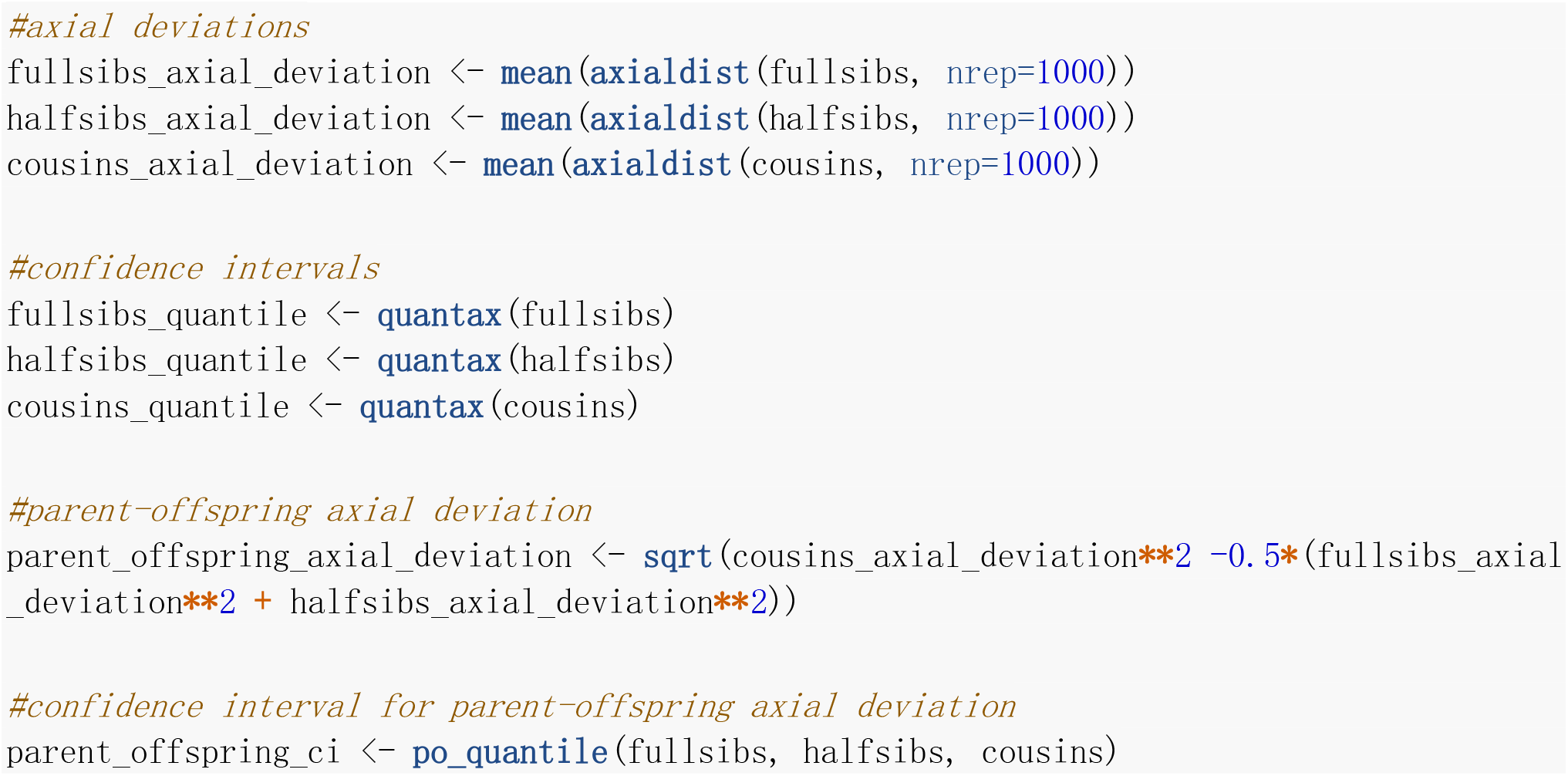

## References

Bravington, M. V., Skaug, H. J., & Anderson, E. C. (2016). Close-kin mark-recapture. Statistical Science, 31(2), 259–274.

Carvalho, D. O., McKemey, A. R., Garziera, L., Lacroix, R., Donnelly, C. A., Alphey, L., … Capurro, M. L. (2015). Suppression of a field population of *Aedes aegypti* in Brazil by sustained release of transgenic male mosquitoes. PLOS Neglected Tropical Diseases, 9(7), e0003864.

Catchen, J., Hohenlohe, P. A., Bassham, S., Amores, A., & Cresko, W. A. (2013). Stacks: an analysis tool set for population genomics. Molecular Ecology, 22(11), 3124–3140.

Cho, K., & Dupuis, J. (2009). Handling linkage disequilibrium in qualitative trait linkage analysis using dense SNPs: a two-step strategy. BMC Genetics, 10(1), 44.

Christophers, S. (1960). Aedes aegypti (L.) the Yellow Fever Mosquito: its Life History, Bionomics and Structure. London: Cambridge University Press.

Clobert, J., Baguette, M., Benton, T. G., & Bullock, J. M. (2012). Dispersal Ecology and Evolution: Oxford University Press.

Danecek, P., Auton, A., Abecasis, G., Albers, C. A., Banks, E., DePristo, M. A., … Group, G. P. A. (2011). The variant call format and VCFtools. Bioinformatics, 27(15), 2156–2158.

Dudchenko, O., Batra, S. S., Omer, A. D., Nyquist, S. K., Hoeger, M., Durand, N. C., … Aiden, A. P. (2017). De novo assembly of the *Aedes aegypti* genome using Hi-C yields chromosome-length scaffolds. Science, 356(6333), 92–95.

Edman, J. D., Scott, T. W., Costero, A., Morrison, A. C., Harrington, L. C., & Clark, G. G. (1998). *Aedes aegypti* (Diptera: Culicidae) movement influenced by availability of oviposition sites. Journal of Medical Entomology, 35(4), 578–583.

Goldberg, C. S., & Waits, L. P. (2010). Quantification and reduction of bias from sampling larvae to infer population and landscape genetic structure. Molecular Ecology Resources, 10(2), 304–313.

Hardy, O. J., & Vekemans, X. (2002). Spagedi: a versatile computer program to analyse spatial genetic structure at the individual or population levels. Molecular Ecology Notes, 2(4), 618–620.

Harrington, L. C., Scott, T. W., Lerdthusnee, K., Coleman, R. C., Costero, A., Clark, G. G., … Edman, J. D. (2005). Dispersal of the dengue vector *Aedes aegypti* within and between rural communities. The American Journal of Tropical Medicine and Hygiene, 72(2), 209–220.

Hauser, L., Baird, M., Hilborn, R., Seeb, L. W., & Seeb, J. E. (2011). An empirical comparison of SNPs and microsatellites for parentage and kinship assignment in a wild sockeye salmon (*Oncorhynchus nerka*) population. Molecular Ecology Resources, 11, 150–161.

Hedrick, P. W., & Kalinowski, S. T. (2000). Inbreeding depression in conservation biology. Annual Review of Ecology and Systematics, 31(1), 139–162.

Hemme, R. R., Thomas, C. L., Chadee, D. D., & Severson, D. W. (2010). Influence of urban landscapes on population dynamics in a short-distance migrant mosquito: evidence for the dengue vector *Aedes aegypti*. PLOS Neglected Tropical Diseases, 4.

Hoffmann, A. A., Iturbe-Ormaetxe, I., Callahan, A. G., Phillips, B. L., Billington, K., & Axford, J. K. (2014). Stability of the *wMel Wolbachia* Infection following invasion into *Aedes aegypti* populations. PLOS Neglected Tropical Diseases, 8(9), e3115.

Hoffmann, A. A., Montgomery, B. L., Popovici, J., Iturbe-Ormaetxe, I., Johnson, P. H., Muzzi, F., … O’Neill, S. L. (2011). Successful establishment of *Wolbachia* in *Aedes* populations to suppress dengue transmission. Nature, 476(7361), 454.

Honorio, N. A., Silva Wda, C., Leite, P. J., Goncalves, J. M., Lounibos, L. P., & Lourenco-de-Oliveira, R. (2003). Dispersal of *Aedes aegypti* and *Aedes albopictus* (Diptera: Culicidae) in an urban endemic dengue area in the State of Rio de Janeiro, Brazil. Memórias do Instituto Oswaldo Cruz, 98(2), 191–198.

Kalinowski, S. T., Wagner, A. P., & Taper, M. L. (2006). ML-RELATE: a computer program for maximum likelihood estimation of relatedness and relationship. Molecular Ecology Notes, 6(2), 576–579.

Kaur, J. S., Lai, Y. L., & Giger, A. D. (2003). Learning and memory in the mosquito *Aedes aegypti* shown by conditioning against oviposition deterrence. Medical and Veterinary Entomology, 17(4), 457–460.

Killeen, G. F., Knols, B. G. J., & Gu, W. (2003). Taking malaria transmission out of the bottle: implications of mosquito dispersal for vector-control interventions. The Lancet Infectious Diseases, 3(5), 297–303.

Langmead, B., & Salzberg, S. L. (2012). Fast gapped-read alignment with Bowtie 2. Nature Methods, 9(4), 357–359.

Loiselle, B. A., Sork, V. L., Nason, J., & Graham, C. (1995). Spatial genetic-structure of a tropical understory shrub, *Psychotria officinalis* (*Rubiaceae*). American Journal of Botany, 82(11), 1420–1425.

Lounibos, L. P. (2003). Genetic-control trials and the ecology of *Aedes aegypti* at the Kenya coast. In W. Takken & T. W. Scott (Eds.), Ecological Aspects for Application of Genetically Modified Mosquitoes (pp. 33–46). Dordrecht: Kluwer Academic Press.

Luikart, G., Ryman, N., Tallmon, D. A., Schwartz, M. K., & Allendorf, F. W. (2010). Estimation of census and effective population sizes: the increasing usefulness of DNA-based approaches. Conservation Genetics, 11(2), 355–373.

Maciel-de-Freitas, R., Codeco, C. T., & Lourenco-de-Oliveira, R. (2007). Daily survival rates and dispersal of *Aedes aegypti* females in Rio de Janeiro, Brazil. The American Journal of Tropical Medicine and Hygiene, 76(4), 659–665.

Maino, J. L., Binns, M., & Umina, P. (2018). No longer a west-side story–pesticide resistance discovered in the eastern range of a major Australian crop pest, *Halotydeus destructor* (Acari: Penthaleidae). Crop and Pasture Science, 69(2), 216–221.

Morrison, A. C., Zielinski-Gutierrez, E., Scott, T. W., & Rosenberg, R. (2008). Defining Challenges and Proposing Solutions for Control of the Virus Vector Aedes aegypti. PLOS Medicine, 5(3), e68.

Nunney, L. (2016). The effect of neighborhood size on effective population size in theory and in practice. Heredity, 117, 224.

O’Neill, S. L., Ryan, P. A., Turley, A. P., Wilson, G., Retzki, K., Iturbe-Ormaetxe, I., … Ritchie, S. A. (2018). Scaled deployment of *Wolbachia* to protect the community from dengue and other *Aedes* transmitted arboviruses. Gates Open Research, 2.

Oksanen, J., Blanchet, F. G., Kindt, R., Legendre, P., O’hara, R., Simpson, G. L., … Wagner, H. (2010). vegan: Community Ecology Package. R package version 1.17-2.

Ordonez-Gonzalez, J. G., Mercado-Hernandez, R., Flores-Suarez, A. E., & Fernandez-Salas, I. (2001). The use of sticky ovitraps to estimate dispersal of *Aedes aegypti* in northeastern Mexico. Journal of the American Mosquito Control Association, 17(2), 93–97.

Ouborg, N. J., Piquot, Y., & Van Groenendael, J. M. (1999). Population genetics, molecular markers and the study of dispersal in plants. Journal of Ecology, 87(4), 551–568.

Palsbøll, P., Zachariah, M. P., & Bérubé, M. (2010). Detecting populations in the ‘ambiguous’ zone: kinship-based estimation of population structure at low genetic divergence. Molecular Ecology Resources, 10(5), 797–805.

Palstra, F. P., & Fraser, D. J. (2012). Effective/census population size ratio estimation: a compendium and appraisal. Ecology and Evolution, 2(9), 2357–2365.

Phillips, C., García-Magariños, M., Salas, A., Carracedo, A., & Lareu, M. V. (2012). SNPs as Supplements in Simple Kinship Analysis or as Core Markers in Distant Pairwise Relationship Tests: When Do SNPs Add Value or Replace Well-Established and Powerful STR Tests? Transfusion Medicine and Hemotherapy: Offizielles Organ der Deutschen Gesellschaft fur Transfusionsmedizin und Immunhamatologie, 39(3), 202–210.

Powell, J. R., & Tabachnick, W. J. (2013). History of domestication and spread of *Aedes aegypti*--a review. Memórias do Instituto Oswaldo Cruz, 108 Suppl 1, 11–17.

Rašić, G., Filipović, I., Weeks, A. R., & Hoffmann, A. A. (2014). Genome-wide SNPs lead to strong signals of geographic structure and relatedness patterns in the major arbovirus vector, *Aedes aegypti*. BMC Genomics, 15, 275.

Reiter, P. (2007). Oviposition, dispersal, and survival in *Aedes aegypti:* implications for the efficacy of control strategies. Vector-Borne and Zoonotic diseases, 7(2), 261–273.

Reiter, P., Amador, M. A., Anderson, A. R., & Clark, G. G. (1995). Short Report: Dispersal of *Aedes aegypti* in an urban area after blood feeding as demonstrated by rubidium-marked eggs. The American Journal of Tropical Medicine and Hygiene, 52(2), 177–179.

Rousset, F. (2000). Genetic differentiation between individuals. Journal of Evolutionary Biology, 13, 58–62.

Russell, R. C., Webb, C. E., Williams, C. R., & Ritchie, S. A. (2005). Mark-release-recapture study to measure dispersal of the mosquito *Aedes aegypti* in Cairns, Queensland, Australia. Medical and Veterinary Entomology, 19(4), 451–457.

Saarman, N. P., Gloria-Soria, A., Anderson, E. C., Evans, B. R., Pless, E., Cosme, L. V., … Powell, J. R. (2017). Effective population sizes of a major vector of human diseases, *Aedes aegypti*. Evolutionary Applications, 10(10), 1031–1039.

Sahlsten, J., Thorngren, H., & Hoglund, J. (2008). Inference of hazel grouse population structure using multilocus data: a landscape genetic approach. Heredity (Edinb), 101(6), 475–482.

Schmidt, T. L., Barton, N. H., Rašić, G., Turley, A. P., Montgomery, B. L., Iturbe-Ormaetxe, I., … Turelli, M. (2017). Local introduction and heterogeneous spatial spread of dengue-suppressing *Wolbachia* through an urban population of *Aedes aegypti*. PLOS Biology, 15(5), e2001894.

Schmidt, T. L., Filipović, I., Hoffmann, A. A., & Rašić, G. (2018). Fine-scale landscape genomics helps explain the slow spatial spread of *Wolbachia* through the *Aedes aegypti* population in Cairns, Australia. Heredity (Edinb), 120(5), 386.

Schmidt, T. L., Rašić, G., Zhang, D., Zheng, X., Xi, Z., & Hoffmann, A. A. (2017). Genome-wide SNPs reveal the drivers of gene flow in an urban population of the Asian Tiger Mosquito, *Aedes albopictus*. PLOS Neglected Tropical Diseases, 11(10), e0006009.

Schultz, A. J., Cristescu, R. H., Littleford-Colquhoun, B. L., Jaccoud, D., & Frère, C. H. (2018). Fresh is best: Accurate SNP genotyping from koala scats. Ecology and Evolution, 8(6), 3139–3151.

Sheppard, P., Macdonald, W., Tonn, R., & Grab, B. (1969). The dynamics of an adult population of *Aedes aegypti* in relation to dengue haemorrhagic fever in Bangkok. The Journal of Animal Ecology, 661–702.

Skrbinšek, T., Jelenčič, M., Waits, L., Kos, I., Jerina, K., & Trontelj, P. (2012). Monitoring the effective population size of a brown bear (*Ursus arctos*) population using new single-sample approaches. Molecular Ecology, 21(4), 862–875.

Szczys, P., Oswald, S. A., & Arnold, J. M. (2017). Conservation implications of long-distance migration routes: Regional metapopulation structure, asymmetrical dispersal, and population declines. Biological Conservation, 209, 263–272.

Turelli, M., & Barton, N. H. (2017). Deploying dengue-suppressing *Wolbachia*: Robust models predict slow but effective spatial spread in *Aedes aegypti*. Theoretical Population Biology, 115, 45–60.

Watts, P. C., Rousset, F., Saccheri, I. J., Leblois, R., Kemp, S. J., & Thompson, D. J. (2007). Compatible genetic and ecological estimates of dispersal rates in insect (*Coenagrion mercuriale:* Odonata: Zygoptera) populations: analysis of ‘neighbourhood size’ using a more precise estimator. Molecular Ecology, 16(4), 737–751.

Wright, S. (1922). Coefficients of inbreeding and relationship. The American Naturalist, 56(645), 330–338.

Wright, S. (1946). Isolation by distance under diverse systems of mating. Genetics, 31(1), 39–59.

